# The Collaborative Cross strains and their founders vary widely in cocaine-induced behavioral sensitization

**DOI:** 10.1101/2022.02.01.478694

**Authors:** SA Schoenrock, L Gagnon, A Olson, M Leonardo, VM Phillip, H He, JD Jentsch, EJ Chesler, LM Tarantino

## Abstract

Cocaine use and overdose deaths attributed to cocaine have increased significantly in the United States in the last 10 years. Despite the prevalence of cocaine use disorder (CUD) and the personal and societal problems it presents, there are currently no approved pharmaceutical treatments. The absence of treatment options is due, in part, to our lack of knowledge about the etiology of CUDs. There is ample evidence that genetics plays a role in increasing CUD risk but thus far, very few risk genes have been identified in human studies. Genetic studies in mice have been extremely useful for identifying genetic loci and genes, but have been limited to very few genetic backgrounds, leaving substantial phenotypic and genetic diversity unexplored.

Herein we report the measurement of cocaine-induced behavioral sensitization using a 19-day protocol that captures baseline locomotor activity, acute locomotor response to cocaine and locomotor sensitization across 5 exposures to the drug. These behaviors were measured in 51 genetically diverse yet tractable Collaborative Cross (CC) strains along with their inbred founder strains. The CC was generated by crossing 8 genetically diverse inbred strains such that each inbred CC strain has genetic contributions from each of the founder strains. Inbred CC mice are infinitely reproducible and provide a stable, yet diverse genetic platform on which to study the genetic architecture and genetic correlations among phenotypes.

We have identified significant differences in cocaine locomotor sensitivity and behavioral sensitization across the panel of CC strains and their founders. We have established relationships among cocaine sensitization behaviors and identified extreme responding strains that can be used in future studies aimed at understanding the genetic, biological and pharmacological mechanisms that drive addiction-related behaviors. Finally, we have determined that these behaviors exhibit relatively robust heritability making them amenable to future genetic mapping studies to identify addiction risk genes and genetic pathways that can be studied as potential targets for the development of novel therapeutics.

## 1. Introduction

Cocaine remains one of the most used illicit substances worldwide and use is particularly high in the United States [1, 2]. Cocaine use in the US has increased significantly over the last 10 years as have overdose deaths attributed to the drug [3, 4]. The personal and societal burdens imposed by cocaine use disorders (CUDs) are staggering and relatively few effective treatment options exist. Not all who use cocaine will develop a CUD, suggesting that there are individual differences in risk. Identifying and understanding the mechanisms that contribute to risk will drive development of novel treatments and prevention strategies.

Twin studies and genome-wide association studies in humans support a significant role for genetic factors in CUD risk [5–7]. To date, however, very few causal genes have been identified in human studies. Animal models have provided important insights into the genetic and neurobiological pathways associated with drug response and reward [8–12]. These experimental systems are a valuable resource that allow for examination of drug effects from the first exposure in naïve animals through more extended exposure paradigms that model the rewarding, reinforcing and motivational properties of drugs. Intravenous drug self-administration (IVSA) protocols are generally considered the “gold standard” for modeling addiction in rodent models. However, IVSA is technically challenging in mice, is labor- and time-intensive and often difficult to establish. Since almost all drugs of abuse increase locomotor activity and there is considerable overlap in dopaminergic mechanisms involved in both psychostimulant-induced locomotor activation and drug reward pathways [13], drug-induced locomotor activation protocols have been used extensively in rodent addiction research.

Acute locomotor activation upon initial drug exposure is frequently used as a model of initial drug sensitivity. In humans, initial sensitivity predicts future drug use but a positive association between initial sensitivity and the development of a substance use disorder has not been well-established [14]. Augmented locomotor activation observed upon repeated exposures, termed behavioral sensitization, is frequently used in model organisms as an indirect measure of underlying neural adaptations that occur in response to chronic drug use [13, 15–19]. Previous work has established two distinct components or phases of behavioral sensitization at the pharmacological and neuroanatomical levels: initiation/development and expression/maintenance of the sensitized behavioral response [16, 20, 21]. The neural adaptations that occur upon repeated exposure to the drug across these phases and the behavioral changes that result have been shown to persist long after withdrawal and are thought to underly continued and persistent drug-taking and -seeking behavior in addicted individuals leading to a high risk of relapse [13, 19, 22]. Although not universally observed, numerous studies have reported a positive relationship between behavioral sensitization and IVSA. Repeated exposure to sensitizing doses of cocaine enhances acquisition of drug self-administration [23, 24] and, conversely, repeated cocaine exposure during self-administration results in sensitized locomotor response to the drug [25–28]. Although the relationship between these behaviors is likely complex and will vary across different populations and genetic backgrounds, these studies support the use of locomotor response to acute and repeated drug exposures as an accessible paradigm for studying genetic and neurobiological mechanisms that influence drug reward and response. In fact, studying behavioral sensitization in diverse genetic backgrounds can reveal relationships among addiction-relevant behaviors that have, as yet, gone undiscovered.

Genetic loci that influence cocaine-induced acute locomotor sensitivity and behavioral sensitization have been identified in numerous studies. However, these behaviors have been examined in very few genetic backgrounds, limiting the general applicability and translatability of the findings [29–38]. The Collaborative Cross (CC) is a panel of recombinant inbred lines derived from intercrossing 5 classical and 3 wild-derived inbred mouse strains that, collectively, capture almost 90% of the known genetic diversity in laboratory mice [39]. Surveying the phenotypic diversity present in the CC offers the opportunity to explore complex behaviors on a genetically defined and reproducible population of inbred mouse strains that have been optimized for systems genetics studies.

We assessed cocaine-induced acute locomotor sensitivity and behavioral sensitization in a set of 51 CC strains and their 8 inbred founders. These genetic reference populations have distinct advantages for the investigation of complex traits. Both inbred populations are ideal for investigating trait correlations, establishing the genetic architecture of quantitative traits and estimating trait heritabilities. Moreover, phenotypic investigation of a panel of CC mice can yield strains with extreme or abnormal phenotypes that can serve as disease models for mechanistic and pharmacologic studies[40]. In this study, we also derived discrete phenotypic variables that will be used for future genetic mapping and gene expression studies in the Diversity Outbred (DO) mouse population. The DO population is derived from the same 8 CC founder strains and is maintained as an outbred population that is ideally suited to gene co-expression analyses and genetic mapping studies [41].

Our data highlight the significant phenotypic variation present for acute and sensitized cocaine-induced locomotor response in this genetically heterogeneous population. The availability of these data in the CC and founder strains affords the opportunity to assess the relationship of these behaviors with other phenotypes as more data are collected and made public in this valuable genetic resource.

## 2 Materials and Methods

### 2.1 Mice

Cohorts of male and female A/J, C57BL/6J, 129S1/SvlmJ, NOD/ShiLtJ, NZO/HlLtJ, CAST/EiJ, PWK/PhJ, WSB/EiJ and CC strains (see **Supplemental Table 1**) were purchased from The Jackson Laboratory (JAX) for behavioral testing in the Center for Systems Neurogenetics of Addiction (CSNA) Behavioral Phenotyping Core. Mice at JAX were housed in specific pathogen-free facilities on a 12-hr light/dark cycle with lights on at 6:00 A.M. Food (NIH31 5K52 chow, LabDiet/PMI Nutrition, St. Louis, MO) and water were provided *ad libitum* throughout the experiment. Animals were singly housed starting at 6 weeks of age until the completion of behavioral testing. Prior to behavioral sensitization, mice were tested in a four-day battery that included open field, light/dark, holeboard and novelty place preference assays (data and methods described in Saul et al 2020 [42] and will not be discussed here). Detailed standard operating procedures for these behavioral assays can be found at https://www.jax.org/research-and-faculty/research-centers/systems-neurogenetics/data-resources. Mice were moved to an anteroom for acclimation for a minimum of 30 minutes prior to behavioral testing by a team of animal handlers; testing by each handler was randomized across batches and strains. Mice tested at JAX averaged 78 days of age at the time of testing (range 49 – 143 days).

Separate cohorts of male and female A/J, C57BL/6J, 129S1/SvlmJ, NOD/ShiLtJ, NZO/HlLtJ, CAST/EiJ, PWK/PhJ, WSB/EiJ mouse strains were purchased from the Jackson Laboratory and transported to the University of North Carolina (UNC) for behavioral sensitization testing. Male and female CC016/GeniUncJ, CC061/GeniUncJ and CC074/UncJ mice were purchased from the UNC Systems Genetics Core Facility. All mice were group housed in a specific pathogen-free facility at UNC and maintained on a 12-hr light/dark cycle with lights on at 7:00 A.M. Food (Harlan Teklad 2920, Envigo, Frederick, MD, USA) and water were provided *ad libitum* throughout the experiment. Mice tested at UNC were behaviorally naïve prior to behavioral sensitization tests and were transported directly from the animal holding room to the procedure space immediately prior to testing. All mice at UNC were tested by the same animal handler. Mice tested at UNC averaged 75 days of age at the time of testing (range 70 – 100 days).

All procedures conducted at JAX and UNC were approved by the Institutional Animal Care and Use Committees at each respective institution and followed guidelines set forth by the National Institutes of Health Guide for the Care and Use of Laboratory Animals, 8^th^ Edition.

### 2.2 Drugs

Cocaine was prepared fresh each test day at a volume 1 mg/mL by dissolving cocaine hydrochloride (Sigma-Aldrich, St. Louis, MO) in 0.9% saline. Cocaine was administered by intraperitoneal (i.p.) injection at a dose of 10 mg/kg in a volume of 0.1 mL/10 g of body weight. Saline injections were also administered by i.p. injection at the same volume.

### 2.3 Open Field Apparatus

The open field arenas were 43.2×43.2×33 cm with white Plexiglas floors and clear Plexiglas walls (ENV-515-16, Med Associates, St. Albans, VT, USA). Infrared photobeam sensors at 2.54 cm intervals on the x, y, and z-axes tracked the animals’ position and activity automatically during testing. Each OF was enclosed in a sound-attenuating chamber (73.5 x 59 x 59 cm) illuminated by two overhead light fixtures containing 28-V lamps.

### 2.4 Behavioral Sensitization to Cocaine

Behavioral sensitization to cocaine was tested in the open field arena (described above). See **Table 1** for a description of test days and treatment groups. Mice were tested for 90 min each test day in a 19-day protocol. Mice were placed into the open field for 30 min, removed and injected intraperitoneally with either saline or cocaine and returned to the arena for 60 min. The CC and founder strain surveys included two test groups – cocaine-treated and saline-treated. Mice in the cocaine-treated group received saline injections on Days 1, 2 and 12 and cocaine injections (10 mg/kg) on Days 3, 5, 7, 9, 11 and 19. All other days were non-testing days and mice were not handled on those days. Mice in the saline group received only saline injections on each test day. Total distance moved in the 60-mins post injection was used for all data analyses and to derive behavioral measures outlined in **Table 2**.

**Table 1.** Behavioral sensitization protocol.

| DAY | 1 | 2 | 3 | 4 | 5 | 6 | 7 | 8 | 9 | 10 | 11 | 12 | 19 |
| --- | --- | --- | --- | --- | --- | --- | --- | --- | --- | --- | --- | --- | --- |
| Drug-Exposed Group | SAL | SAL | COC | No Testing | COC | No Testing | COC | No Testing | COC | No Testing | COC | SAL | COC |
| Saline-Exposed Group | SAL | SAL | SAL |  | SAL |  | SAL |  | SAL |  | SAL | SAL | SAL |

SAL = Saline, COC = Cocaine

**Table 2.** Derived variables calculated from 19-day behavioral sensitization data.

| BEHAVIORAL VARIABLE | DESCRIPTION (DERIVED VARIABLE) |
| --- | --- |
| Day 1 Locomotion | Distance traveled on 1st exposure to open field and saline injection (Day 1) |
| Habituated Locomotion | Distance traveled on 2nd exposure to open field and saline injection (Day 2) |
| Acute Locomotor Response | Locomotor response after 1st exposure to cocaine (Day 3 – Day 2) |
| Initial Locomotor Sensitization | Distance traveled after 2nd cocaine exposure (Day 5 – Day 3) |
| Locomotor Sensitization | Area under the curve (AUC) of distance across 5 consecutive cocaine exposures (Day 3,5,7,9,11) |
| Sensitization Expression | Locomotor response to cocaine after 1-week withdrawal (Day 19 – Day 11) |
| Conditioned Activation | Locomotor response to saline after repeated cocaine exposures (Day 12 – Day 2) |

### 2.5 Statistical Analyses

Open field behavioral data were analyzed post-session using commercially available software (Activity Monitor 7.06, Med Associates). All statistical analyses were performed using R Studio 1.2.13335 or SPSS Statistics v26 for Mac OS X 10.6+. Graphs were generated using Graphpad Prism 8 for Mac OS X. Descriptive statistics were examined to show the mean and standard deviation for each strain and each study at different days. A three-way linear mixed effects ANOVA was performed using the R package lmerTest to evaluate the effects of treatment, strain and day on day-specific trait measures with a random effect of mouse. Significant differences involving treatment or strain were further evaluated by post-hoc Tukey’s HSD. Significance for all comparisons was set at α=0.05. ANOVA was also performed to assess replicability of derived variables (**Table 2**) across testing sites. Significant main effects were examined by post-hoc Tukey’s HSD. Spearman correlation analyses were used to assess the relationship among derived variables. Broad sense heritability (H^2^) values were calculated using the cocaine-exposed group using the following equation: [Mean Square Strain Effects/(Mean Square Strain Effect+(Mean Number of Mice per Strain-1)*Mean Square Residuals)].

## 3. Results

### 3.1 Treatment and Strain Effects

A linear mixed model was used to test the effects of test day, strain and treatment (cocaine vs saline) on locomotor activity in CC and founder strains across all 19 days of testing. Significant day, treatment and strain as well as day x treatment, day x strain, treatment x strain and day x treatment x strain interaction effects were observed (all *p*<2.2×10^-16^). Posthoc analyses of treatment effects revealed that 35 of the 58 strains showed a significant increase in locomotor activity when exposed to cocaine vs saline (**Supplemental Table 2**). CC strains that were most significantly activated in response to cocaine include CC004/TauUncJ, the strain identified in our previously published study ([43]; *p*=1.1×10^-27^), as well as CC016/GeniUncJ (*p*=1.3×10^-8^), CC027/GeniUncJ (*p*=4.0×10^-7^) and CC035/UncJ (p=7.6×10^-9^). Several CC strains were also identified as non-responders including CC041/TauUncJ (*p*=0.60) as described in our previously published study [43], as well as CC010/GeniUncJ (*p*=0.97), CC068/TauUncJ (*p*=0.68), CC079/TauUncJ (*p*=0.60) (**Fig 1, Supplemental Fig 1**).

**Figure 1.**
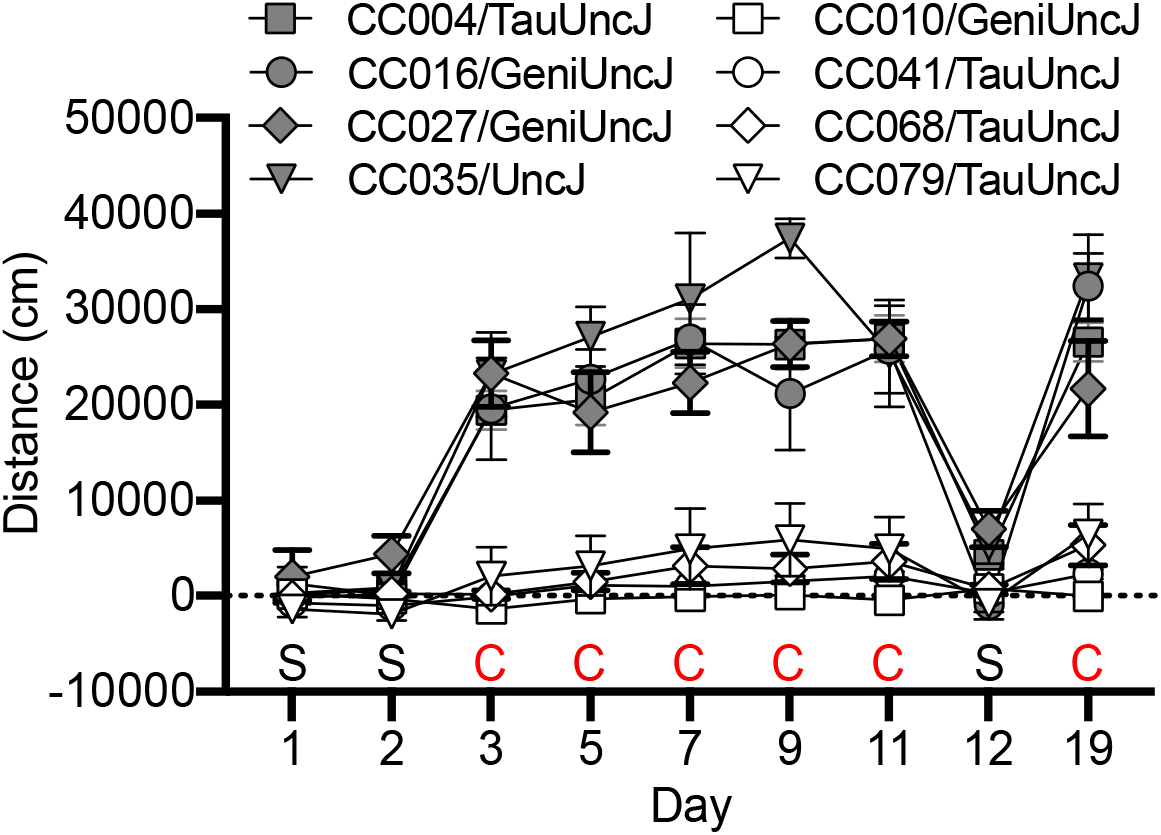
Collaborative Cross and Founder Strains with highest and lowest responses to 10 mg/kg cocaine across the 19-day sensitization protocol. Locomotor activity for each cocaine-exposed mouse was subtracted from the mean of all sham-treated mice in the same strain. Lines depict strain means and error bars are standard error of the mean.

We also assessed treatment and strain effects on derived variables (**Table 2, Supplemental 3**). We observed significant strain effects for Day 1 and Day 2 locomotor activity (both *p*<2×10^-16^), acute locomotor response to cocaine (Day 3 – Day 2; *p*<2×10^-16^), initial sensitization (Day 5 – Day 3; *p*=0.002) and sensitization AUC (*p*=1.1×10^-4^). No significant were observed for sensitization expression (Day 19 – Day 11; *p*=0.061) or conditioned activation (Day 12 – Day 2; *p*=0.38). Significant effects of treatment were observed for all derived variables (all *p*<0.05). There was a significant strain by treatment interaction for acute locomotor response to cocaine (Day 3 – Day 2; *p*<2×10^-16^), initial sensitization (Day 5 – Day 3; *p*=0.025) and sensitization AUC (*p*=0.013).

### 3.2 Sex Effects in the CC Founder Strains

The limited number of male and female mice tested in each CC strain did not allow for a assessment of main effects of sex or interactions. We were able to examine sex differences and strain by sex interactions in the founder strains using a linear mixed model. We observed a significant main effect of sex in the founder group; female mice are more active than male mice (*p* = 0.004). We also observed significant sex by strain effects (*p*=1.4×10^-4^). CAST/EiJ females are significantly more active than CAST/EiJ males (*p*=4.8×10^-3^) as are WSB/EiJ females in comparison to WSB/EiJ males (*p*=2.8×10^-6^). No significant sex differences were observed in the remaining six founder strains (**Fig 2, Supplemental Table 4, Supplemental Fig 2**). We did not identify any significant sex by treatment or sex by day effects.

**Figure 2.**
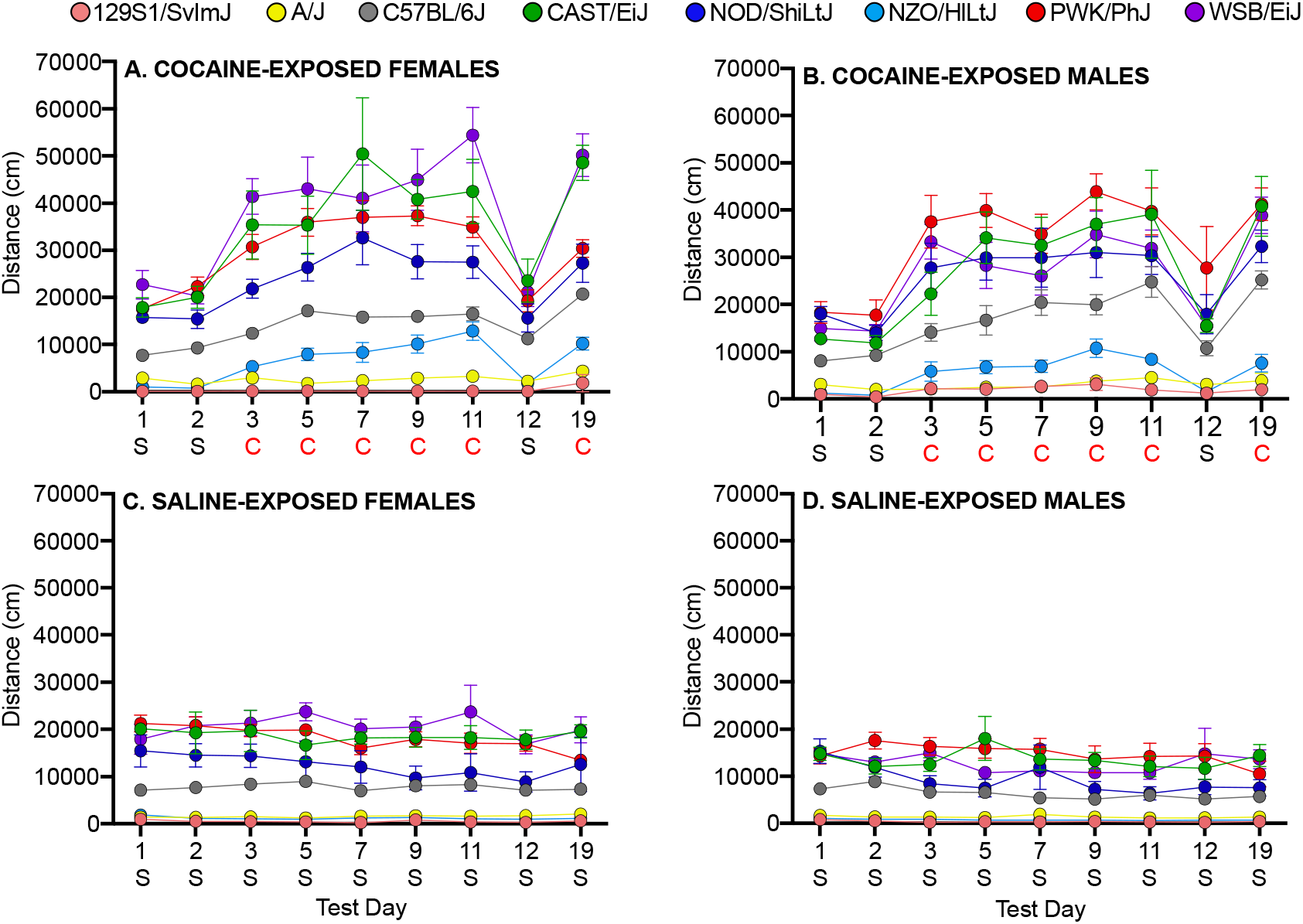
Locomotor activity in response to cocaine and saline in female (**A,C**) and male (**B,D**) founder strain mice. S=Saline, C=Cocaine. Error bars are standard error of the mean.

### 3.3 Heritability

Derived variables were used to generate CC and founder strain means. These data were used to calculate broad sense heritability in the cocaine-treated group to determine the proportion of phenotypic variation due to genetic background. Heritabilities ranged from 0.10 – 0.73 (**Supplemental Table 5A**). The highest heritabilities were observed for locomotor activity on Days 1 and 2 (both H^2^ = 0.73), acute locomotor response to cocaine (Day 3 – Day 2) had a heritability of 0.48 and initial sensitization (Day 5 – Day 3) and behavioral sensitization (AUC) had heritabilities of 0.25 and 0.22, respectively. Sensitization expression (Day 19 – Day 11) and conditioned activation (Day 12 – Day 2) had the lowest heritabilities (H^2^ = 0.16 and 0.10, respectively).

### 3.4 Correlation of Sensitization Behaviors

Correlations between derived variables for all behavioral sensitization behaviors (**Table 2**) were performed using CC and founder strain means for cocaine-exposed groups. We detected significant positive correlations between Day 1 and Day 2 locomotor activity (*p*<0.0001). Day 1 and Day 2 locomotor activity were significantly and positively correlated with acute locomotor response to cocaine (Day 3 – Day 2; *p*<0.0001). Behavioral sensitization (AUC) was significantly and positively correlated with acute locomotor response to cocaine (*p*<0.05), initial sensitization (Day 5 – Day 3; *p*<0.0001) and conditioned activation (Day 12 – Day 2; *p*<0.01). We also observed a significant positive correlation between initial sensitization and conditioned activation (*p*<0.01). (**Fig 3, Supplemental Table 6**)

**Figure 3.**
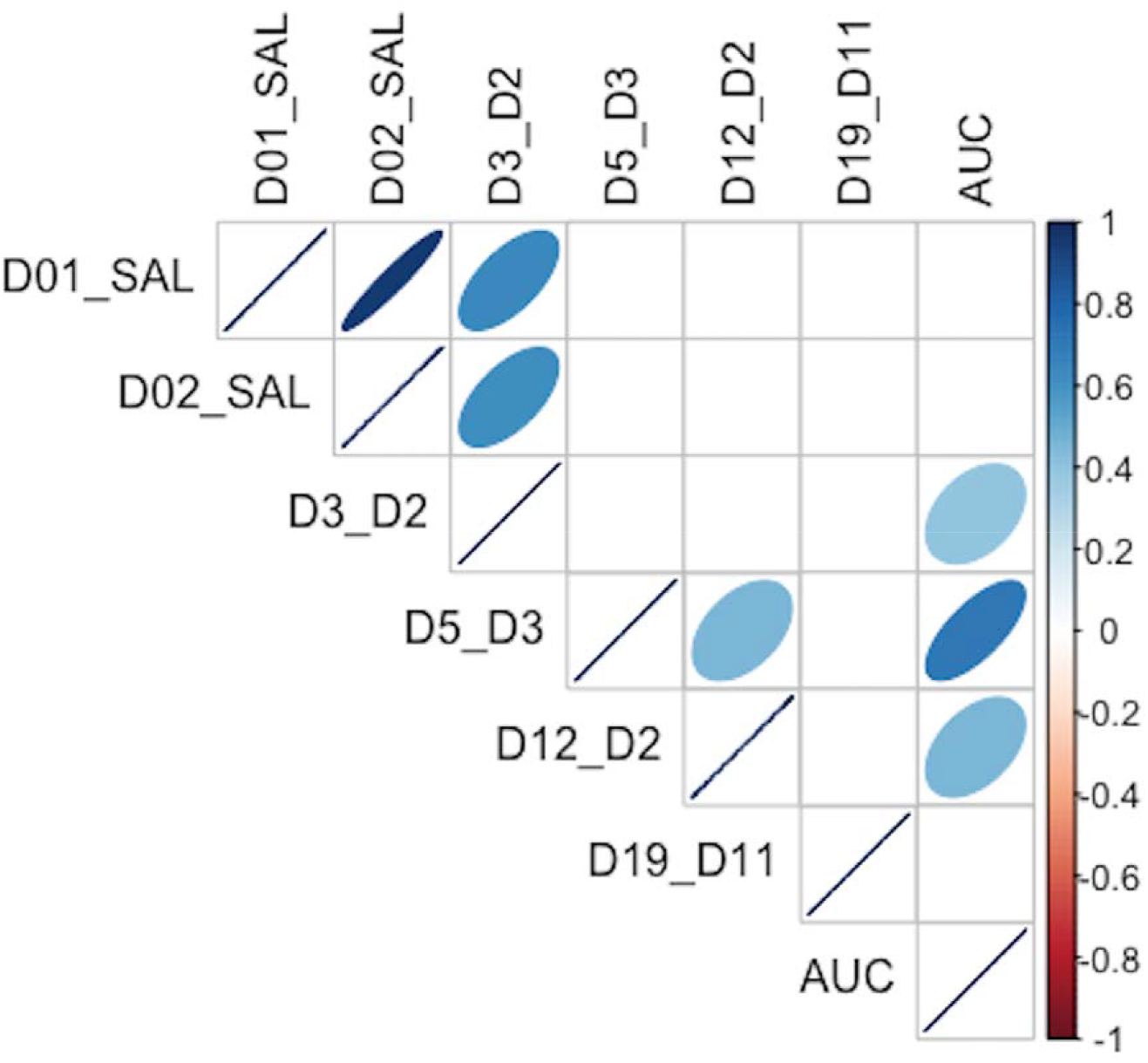
Strain mean correlations among all derived variables in the Collaborative Cross and Founder strains in cocaine-exposed groups. Strength of the correlation is depicted by the color (darker = stronger) and width (narrower = stronger) of the ellipse.

### 3.5 Replicability of Sensitization Behavior

#### 3.5.1 Founder Replicability

A separate cohort of cocaine-exposed founder strain mice were tested at UNC (**Supplemental Table 1**). Comparison of the derived variables measured in the UNC cohort with the cocaine-exposure group of mice tested at JAX allows us to assess the replicability of sensitization behavior across the two sites (**Fig 4**). ANOVA of each behavioral measure with strain, sex and test site as independent factors yielded significant strain effects for Day 1 (F_(7,153)_=69.1; *p*=3.4×10^-44^) and habituated locomotor activity (Day 2; F_(7,151)_=68.9; *p*=7.7×10^-44^), acute locomotor response to cocaine (Day 3 – Day 2; F_(7,151)_=19.0; *p*=4.7×10^-18^), initial sensitization (Day 5 – Day 3; F_(7,150)_=5.8; *p*=6.0×10^-6^) and behavioral sensitization (AUC; F_(7,146)_=10.9; *p*=5.0×10^-11^). We also observed significant sex effects for Day 1 (F_(1,153)_=11.3; *p*=9.7×10^-4^) and habituated locomotor behavior (F_(1,151)_=21.1; *p*=9.0×10^-6^), acute response to cocaine (F_(1,151)_=4.6; *p*=0.034), behavioral sensitization (F_(1,146)_=7.5; *p*=0.007) and conditioned activation (F_(1,149)_=4.9; *p*=0.028). Test site differences were observed for Day 1 (F_(1,153)_=42.7; *p*=8.8×10^-10^) and habituated locomotor behavior (F_(1,153)_=27.0; *p*=6.5×10^-7^), initial sensitization (F_(1,150)_=13.8; *p*=2.9×10^-4^) and behavioral sensitization (F_(1,146)_=16.5; *p*=7.8×10^-5^). Generally, mice tested at JAX were more active than mice tested at UNC. However, significant strain by testing site differences were detected for Day 1 (F_(7,153)_=4.4; *p*=1.7×10^-4^) and habituated locomotor activity (F_(7,151)_=2.5; *p*=0.017) and behavioral sensitization AUC (F_(7,146)_=3.0; *p*=0.005) (**Fig 5A-C**). We also generated Pearson correlation coefficients of derived variables from CC and Founder mice tested at the two sites. Only sensitization expression (Day 19 – Day 11; r(6)=0.34) and conditioned activation (Day 12 – Day 2; r(6)=0.62) were not significantly correlated between sites. The remainder of the derived variables had correlation coefficients ranging from 0.82 – 0.97 (**Fig 4**).

**Figure 4.**
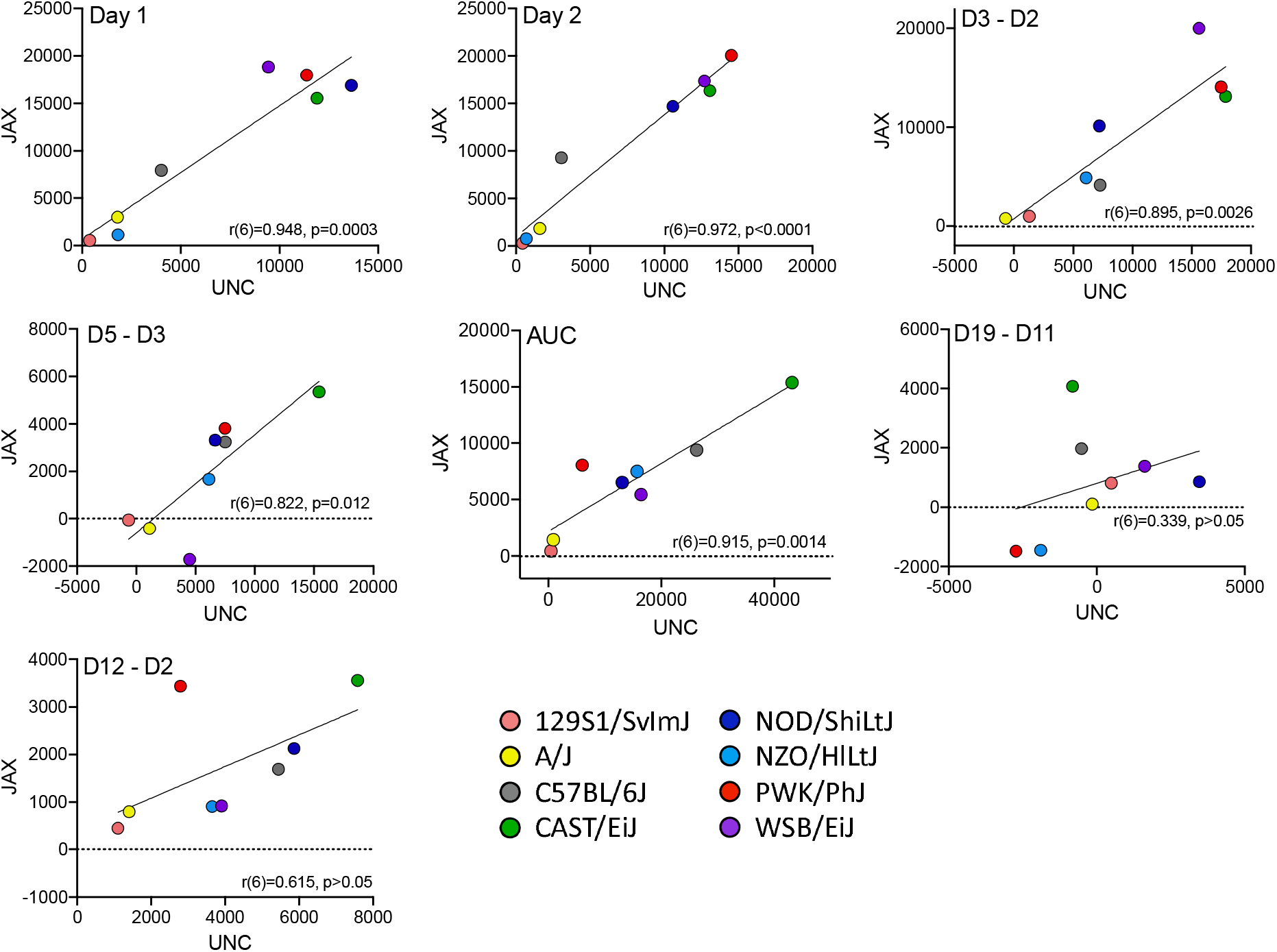
Founder strain means for all derived variables for mice tested at UNC and JAX. Each data point represents a strain mean. The line of best fit and correlation coefficient (r) with *p*-value are shown for each derived variable.

**Figure 5.**
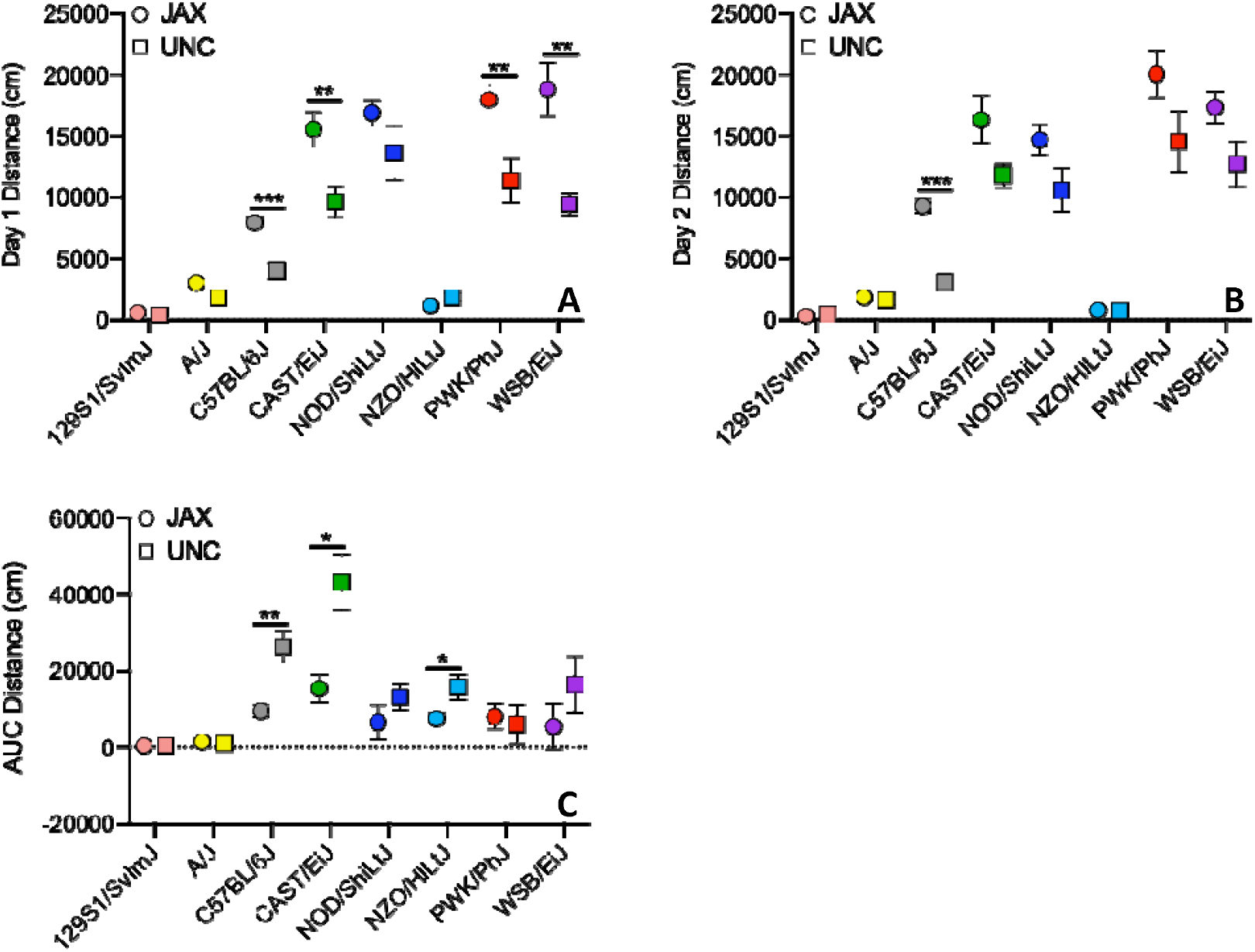
Founder strain means for Day 1 (**A**) and Day 2 distance (**B**) and sensitization AUC (**C**) behaviors measured at JAX (circles) and UNC (squares). Error bars are standard error of the mean. *p<0.05, \*\**p*<0.01, \*\*\**p*<0.001

#### 3.5.2 CC Replicability

In addition to the founder strains, we also tested three CC strains at UNC. We chose two “high responding” strains (CC016/GeniUncJ, CC074/UncJ) and one “low responding” strain (CC061/GeniUncJ) (**Supplemental Table 5**). We analyzed derived sensitization variables using ANOVA with test site, strain and sex as independent factors. We observed significant test site effects for acute response to cocaine (Day 3 – Day 2; F_(1,29)_=7.5; *p*=0.01) and sensitization (AUC; F_(1,29)_=6.98; *p*=0.013). Acute response to cocaine was higher and sensitization AUC was reduced in mice tested at UNC compared to JAX (**Fig 6A,B**). We also observed significant strain by test site interactions for habituated locomotor activity (Day 2; F_(2,28)_=12.8; *p*=1.1×10-4) and conditioned activation (Day 12 – Day 2; F_(2,29)_=3.7; *p*=0.039). Both interaction effects were driven primarily by higher locomotor activity on Day 2 in mice tested at the UNC site (**Fig 6C,D, Supplemental Figure 3**).

**Figure 6.**
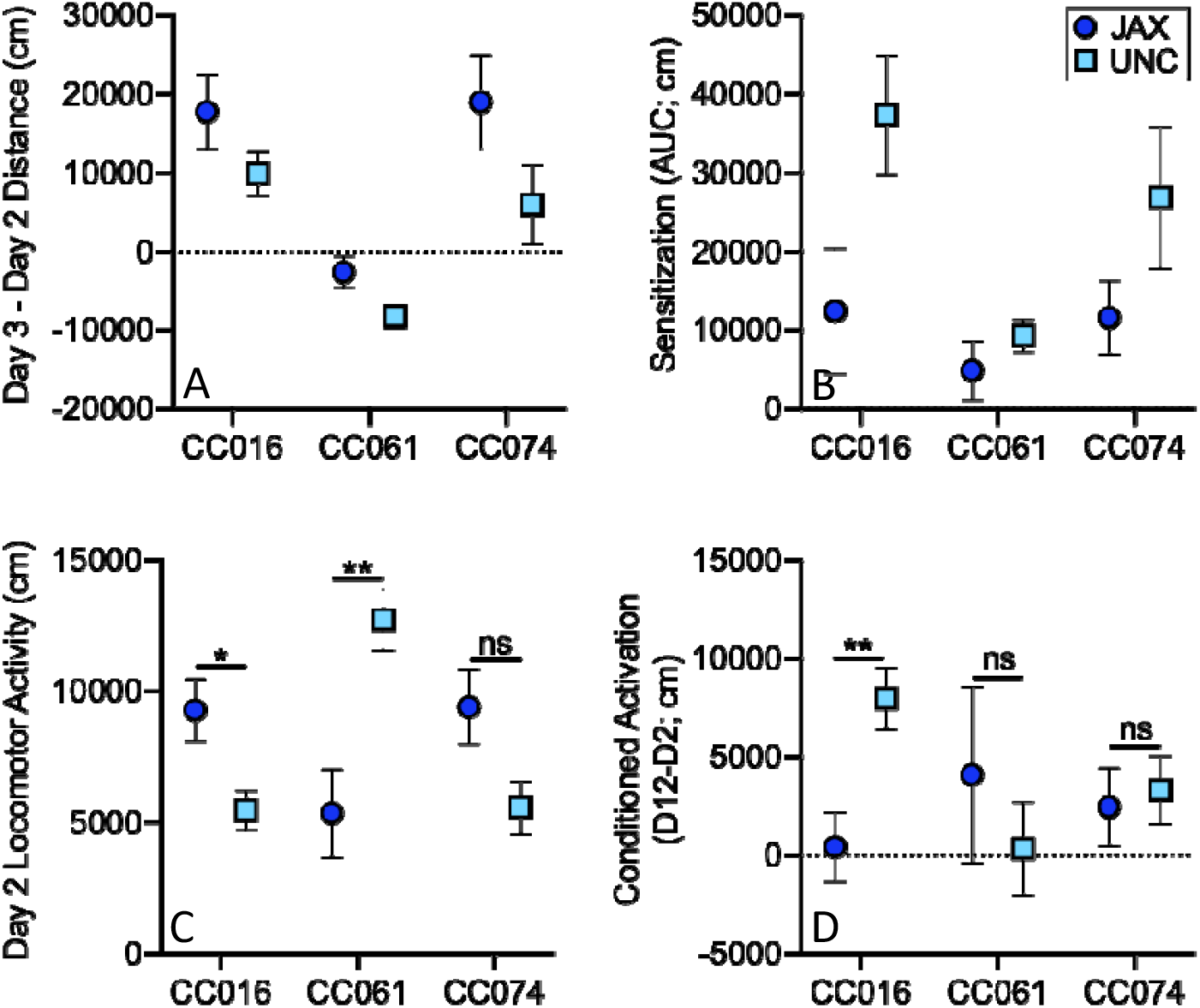
Comparison of CC016/GeniUncJ, CC061/GeniUncJ and CC074/UncJ behavior across test sites. Significant differences in acute locomotor activation (**A**) and sensitization AUC (**B**) were observed across sites as well as strain by test site interactions for habituated locomotor activity (**C**) and conditioned activation (**D**). Each data point is the strain mean and error bars are standard error of the mean. *p<0.05, \*\**p*<0.01, \*\*\**p*<0.001

## 4. Discussion

The absence of effective pharmaceutical treatments for CUD has hindered our ability to effectively combat this devastating disorder. Gaps in our knowledge regarding the underlying etiology of CUD limit our ability to develop novel and efficacious pharmacotherapies. There is strong evidence for genetic factors that influence CUD risk [5–7] but thus far, very few risk genes have been identified in human genome-wide association studies [5, 44, 45]. Preclinical studies in mice have been effective in identifying genetic loci and candidate genes that influence the activating and reinforcing effects of cocaine [30–33, 46], but the majority of these studies have been conducted on only a few inbred strain backgrounds, limiting the scope and generalizability of the findings. In fact, the most commonly used inbred strain, C57BL/6J, falls close to the middle of the phenotypic distribution for many of the sensitization variables that we studied (**Supplemental Figure 1A-G**). The use of the genetically diverse CC population allowed us to capture a broader range of phenotypes that can drive novel biological and mechanistic discoveries. It is clear that the phenotypic distribution of the CC strains exceeds that of the founder strains for almost all behaviors measured with the exception of locomotor activity on Days 1 and 2, measured prior to any drug exposures (**Supplemental Figures 1A-G**).

The ability to identify extreme responding strains is a significant benefit of measuring addiction-like behaviors and other phenotypes in the CC panel. We, and others, have identified extreme CC strains that can be used as models to study the genetic, biological and pharmacological mechanisms of disease [47, 48]. CC004/TauUncJ and CC041/TauUncJ have already been described in our previous publication as CC strains that are behaviorally divergent for the locomotor activating effects of cocaine and also for acquisition of intravenous cocaine self-administration, circadian behavior and tissue content of monoamines [43]. The expanded set of cocaine-induced locomotor sensitization data described here in the CC and founder population provides the opportunity to identify additional inbred CC strains with abnormal or extreme phenotypes to study mechanisms that contribute to cocaine-induced behavioral differences.

Behavioral sensitization occurs in several phases, including locomotor response to an initial exposure to the drug, augmented response upon additional exposures and finally, persistent elevation of the response even after a period of abstinence. Our locomotor sensitization protocol allows us to measure all of these phases and examine them as discrete variables. Examining correlations (or lack thereof) across different behavioral sensitization measures can inform us about genetic and phenotypic relationships across the phases of sensitization and inform future studies aimed at genetic mapping of discrete behaviors. Not surprisingly, the strongest correlation was observed between two behavioral measures of locomotor activity following an injection of saline on Days 1 and 2. These measures were also significantly correlated with acute locomotor response to cocaine (Day 3 – Day 2). The relationship between locomotor activity and psychostimulant-induced locomotor activation has been repeatedly observed in our laboratory [49, 50] and reported by others [51–53]. Neither Day 1 nor Day 2 habituated locomotor response were significantly correlated with any of the behavioral sensitization measures suggesting that locomotor activity itself does not predict the locomotor-sensitizing properties of cocaine in this population. Sensitization expression (Day 19 – Day 11) does not correlate with any other behavioral measure and also has fairly low heritability (H2 = 0.16). Locomotor response to cocaine on Day 19 was remarkably consistent in comparison to Day 11 across all strains even after 7 days of abstinence and regardless of the level of locomotor sensitization across cocaine exposure days (**Supplemental Table 5A**). Our data suggest that in this population, genetic influences are more important in the initiation or development phases and not in the maintenance or expression of the sensitized response. This result impacts our interpretation of the role that genetics may play in cocaine-induced neural adaptations.

The derived sensitization variables (described in **Table 2**) are discrete behavioral measures necessary for cross-strain comparisons and statistical analyses as well as future mapping studies in the outbred DO population. However, these variables do not fully capture the diversity of behavioral responses across the CC and founder strains. Closer examination of behavior across the 19-day sensitization assay can highlight distinct sensitization patterns (as previously described by Smith et al. [54]), that suggest acute differences in drug response (**Fig 7A**), rate of sensitization (**Fig 7B**) and differences in maximal response to cocaine (**Fig 7C**). However, we note that our study was limited to a single dose of cocaine and differences in drug response may be dose-specific. Dose-response studies are necessary prior to moving forward with investigations of the mechanisms that might explain behavioral differences in specific strains of interest.

**Figure 7.**
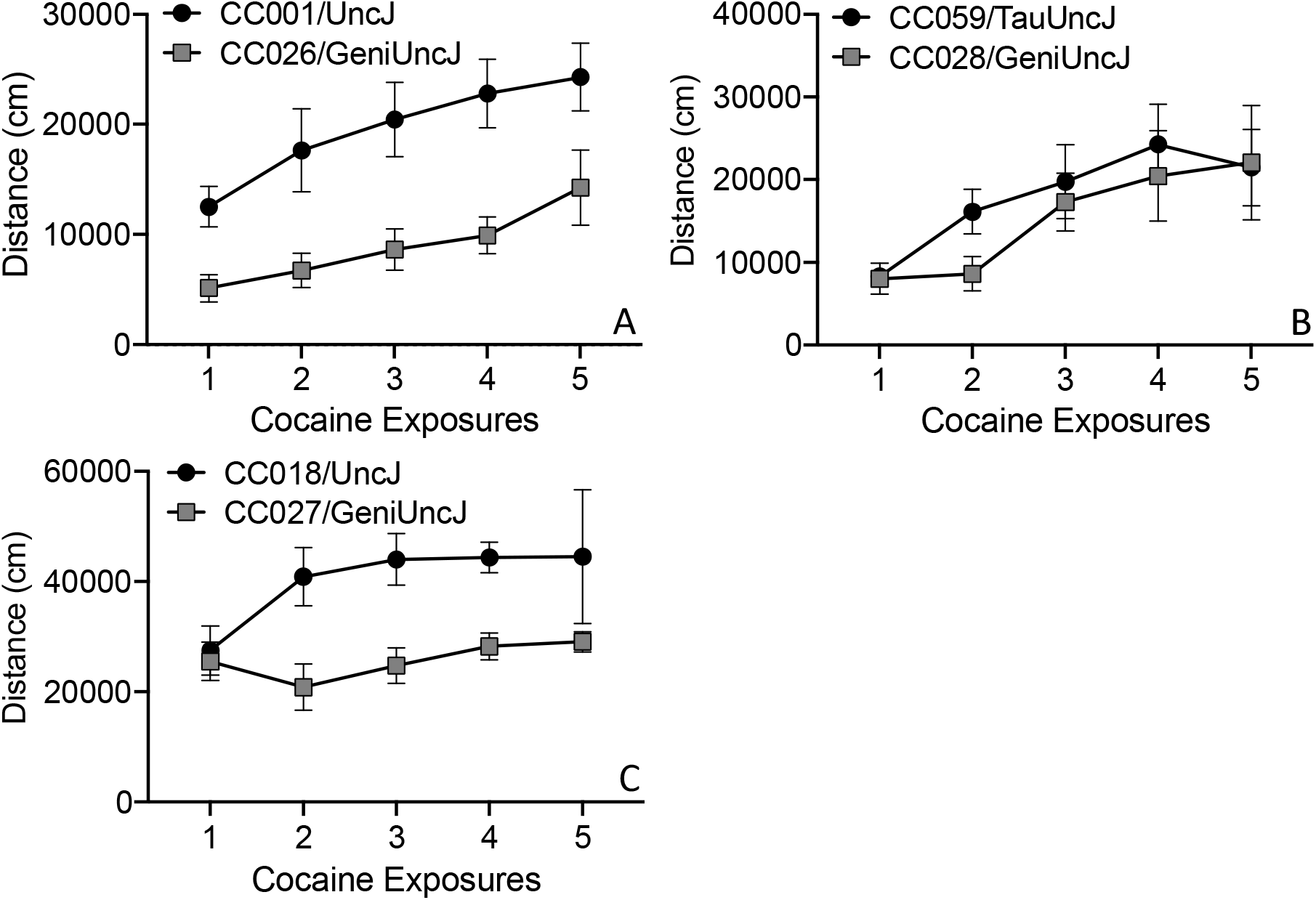
Different patterns of sensitized cocaine locomotor behavior in CC strains of interest. Cocaine exposures 1 to 5 correspond to Days 3 - 11 in the sensitization protocol. (**A**) CC001/UncJ and CC026/GeniUncJ show similar rates of sensitization but differences in acute response; (**B**) CC059/TauUncJ mice exhibit a robust sensization response across the first four exposures while CC028/GeniUncJ mice don’t begin to show a sensitized response until the third exposure and (**C**) the maximal locomotor response in CC018/UncJ mice is reached by the second dose whereas CC027/GeniUncJ mice show the highest response after a single exposure and do not increase upon additional exposures.

Phenotype replicability is an important consideration in preclinical studies and has been discussed in great detail in the field of behavioral genetics [55, 56]. The inability to replicate behaviors across experimental sites, under different protocols or across time can limit the generalizability and translatability of findings and has broader implications for the overall stability and robustness of commonly used behaviors. We did observe differences in behavior in founder strains across sites, but mostly these were main effects that detected similar shifts in behavior across all strains. Strain by test site interactions were observed for a few phenotypes (**Fig 5**) but the correlation between behaviors was remarkably strong between sites (**Fig 4**) even considering differences in testing protocol across sites. A similar outcome was also observed for replication among the CC strains for which we have data from both sites (**Supplemental Figure 3**). Strain by test site interactions mainly appear to be driven by behavioral variability for locomotor activity on Day 2 in CC061 (**Fig 6C**) or for conditioned activation (**Fig 6D**) for which heritability (H^2^ = 0.10) is not particularly robust. These results are encouraging considering the differences in testing protocols at UNC vs JAX (group vs single housing, test naïve mice vs previous exposure to testing, etc) and the procurement of mice from different facilities which introduces additional environmental factors including exposure to different diets during development and post-weaning, transportation etc. Overall, we found that the pattern of cocaine-induced behaviors for a given strain were replicated across test sites. (**Figs 5, 6**).

Our strain survey of cocaine-induced behavioral sensitization in the CC population is a unique resource that will be of great use to the addiction genetics community. Individual strains can be used as disease models for studying neurobiological and pharmacological mechanisms. The heritability observed for most of the sensitization behaviors also indicates that mapping strategies to identify genetic loci and, ultimately, addiction risk genes, will be fruitful. Genetic mapping in the DO population is currently underway in the CSNA. Identification of risk genes in this preclinical model will inform our knowledge of the mechanisms and pathways that affect differential responses to repeated cocaine exposures and generate novel hypotheses regarding the path from initial drug use to a CUD.

