## Supplemental Materials for "The Collaborative Cross strains and their founders vary widely in cocaine-induced behavioral sensitization"

**Supplemental Table 1.** Number of animals tested per strain at both test sites

| Study |  | Primary | | Primary | | Replication Group | |
| --- | --- | --- | --- | --- | --- | --- | --- |
| Testing Site |  | JAX | | JAX | | UNC | |
| Treatment Group |  | Cocaine Exposed | | Saline Only | | Cocaine Exposed | |
| Strain | JAX Stock # | F | M | F | M | F | M |
| 129S1/SvImJ | 002448 | 5 | 6 | 8 | 8 | 4 | 4 |
| A/J | 000646 | 12 | 12 | 6 | 8 | 4 | 4 |
| C57BL/6J | 000664 | 5 | 7 | 7 | 6 | 4 | 8 |
| CAST/EiJ | 000928 | 6 | 5 | 7 | 7 | 4 | 4 |
| NOD/ShiLtJ | 001976 | 6 | 6 | 4 | 5 | 4 | 4 |
| NZO/HlLtJ | 002105 | 6 | 7 | 10 | 9 | 4 | 4 |
| PWK/PhJ | 003715 | 9 | 9 | 8 | 8 | 4 | 4 |
| WSB/EiJ | 001145 | 7 | 7 | 7 | 7 | 4 | 4 |
| CC001/UncJ | 021238 | 4 | 4 | 4 | 4 |  |  |
| CC002/UncJ | 021236 | 2 | 3 | 3 | 2 |  |  |
| CC003/UncJ | 021237 | 2 | 2 | 2 | 2 |  |  |
| CC004/TauUncJ | 020944 | 19 | 21 | 11 | 11 |  |  |
| CC005/TauUncJ | 020945 | 2 | 3 | 4 | 3 |  |  |
| CC006/TauUncJ | 022869 | 2 | 2 | 2 | 2 |  |  |
| CC007/UncJ | 029625 | 4 | 4 | 1 | 2 |  |  |
| CC008/GeniUncJ | 026971 | 5 | 5 | 1 | 1 |  |  |
| CC009/UncJ | 018856 | 2 | 2 | 2 | 2 |  |  |
| CC010/GeniUncJ | 021889 | 2 | 2 | 1 | 2 |  |  |
| CC011/UncJ | 018854 | 5 | 5 | 4 | 5 |  |  |
| CC012/GeniUncJ | 028409 | 5 | 5 | 5 | 5 |  |  |
| CC013/GeniUncJ | 021892 | 4 | 2 | 2 | 3 |  |  |
| CC015/UncJ | 018859 | 4 | 1 | 2 | 1 |  |  |
| CC016/GeniUncJ | 024684 | 4 | 4 | 4 | 4 | 8 | 8 |
| CC017/UncJ | 022870 | 3 | 3 | 3 | 3 |  |  |
| CC018/UncJ | 021890 | 2 | 2 | 2 | 2 |  |  |
| CC019/TauUncJ | 021894 | 4 | 5 | 5 | 5 |  |  |
| CC020/GeniUncJ | 025129 | 2 | 2 | 2 | 2 |  |  |
| CC023/GeniUncJ | 025131 | 2 | 2 | 2 | 2 |  |  |
| CC024/GeniUncJ | 021891 | 1 | 2 | 2 | 2 |  |  |
| CC025/GeniUncJ | 018857 | 2 | 2 | 1 | 2 |  |  |
| CC026/GeniUncJ | 024685 | 4 | 4 | 4 | 4 |  |  |
| CC027/GeniUncJ | 025130 | 3 | 3 | 3 | 3 |  |  |

**Supplemental Table 1 cont’d.** Number of animals tested per strain at both test sites

| Study |  | Primary | | Primary | | Replication Group | |
| --- | --- | --- | --- | --- | --- | --- | --- |
| Testing Site |  | JAX | | JAX | | UNC | |
| Treatment Group |  | Cocaine Exposed | | Saline Only | | Cocaine Exposed | |
| Strain | JAX Stock # | F | M | F | M | F | M |
| CC028/GeniUncJ | 025126 | 4 | 4 | 4 | 4 |  |  |
| CC029/UncJ | 026972 | 2 | 2 | 2 | 2 |  |  |
| CC030/GeniUncJ | 025426 | 3 | 3 | 3 | 3 |  |  |
| CC032/GeniUncJ | 020946 | 3 | 3 | 2 | 1 |  |  |
| CC033/GeniUncJ | 025910 | 2 | 3 | 2 | 1 |  |  |
| CC035/UncJ | 024073 | 3 | 3 | 3 | 2 |  |  |
| CC036/UncJ | 025127 | 3 | 3 | 3 | 3 |  |  |
| CC037/TauUncJ | 025423 | 3 | 3 | 3 | 3 |  |  |
| CC040/TauUncJ | 023831 | 2 | 2 | 2 | 3 |  |  |
| CC041/TauUncJ | 021893 | 20 | 19 | 19 | 20 |  |  |
| CC043/GeniUncJ | 023828 | 2 | 2 | 2 | 2 |  |  |
| CC044/UncJ | 026426 | 2 | 2 | 2 | 1 |  |  |
| CC045/GeniUncJ | 025425 | 2 | 2 | 1 | 1 |  |  |
| CC051/TauUncJ | 021897 | 2 | 3 | 3 | 2 |  |  |
| CC057/UncJ | 024683 | 1 | 1 | 2 | 1 |  |  |
| CC059/TauUncJ | 025125 | 3 | 2 | 2 | 3 |  |  |
| CC060/UncJ | 026427 | 1 | 1 | 1 | 1 |  |  |
| CC061/GeniUncJ | 023826 | 2 | 1 | 1 | 1 | 7 | 6 |
| CC065/UncJ | 023830 | 1 |  | 1 |  |  |  |
| CC068/TauUncJ | 025908 | 2 | 2 | 2 | 2 |  |  |
| CC074/UncJ | 018855 | 2 | 2 | 2 | 2 | 6 | 6 |
| CC075/UncJ | 027293 | 1 | 1 | 1 | 1 |  |  |
| CC078/TauUncJ | 025989 | 2 | 2 | 2 | 2 |  |  |
| CC079/TauUncJ | 025990 | 2 | 2 | 1 | 2 |  |  |
| CC080/TauUncJ | 025988 | 1 | 1 | 2 | 2 |  |  |
| CC083/UncJ | 031921 | 2 | 2 | 1 | 1 |  |  |
| CC084/TauJ | 028923 | 1 | 1 | 1 | 1 |  |  |

**Supplemental Table 2.** *Post hoc* t-tests of cocaine vs saline (SHAM) treatment on locomotor activity in CC and Founder strains

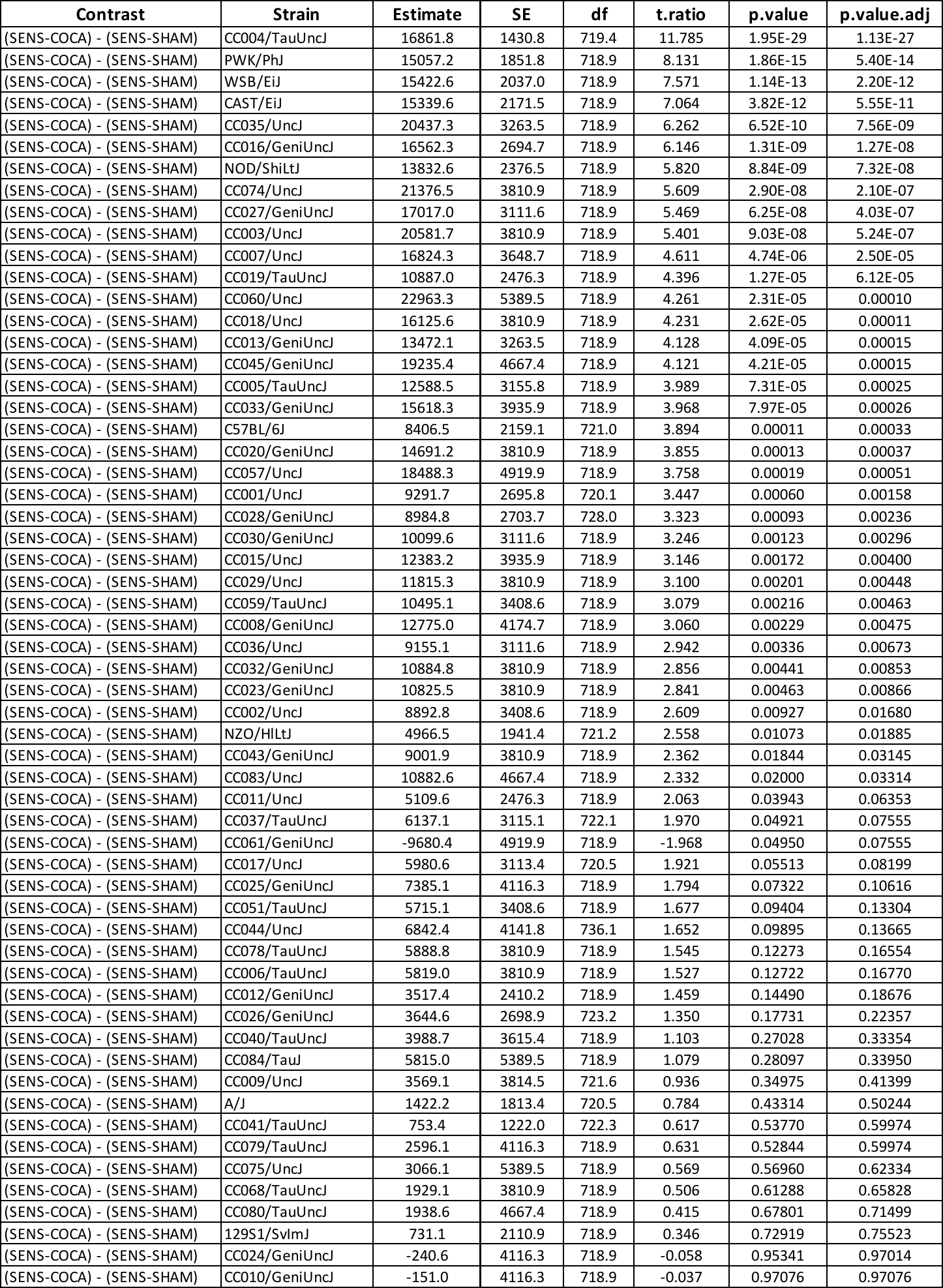

**Supplemental Table 3.** Strain and treatment effects on derived behaviors in the CC and founder strains

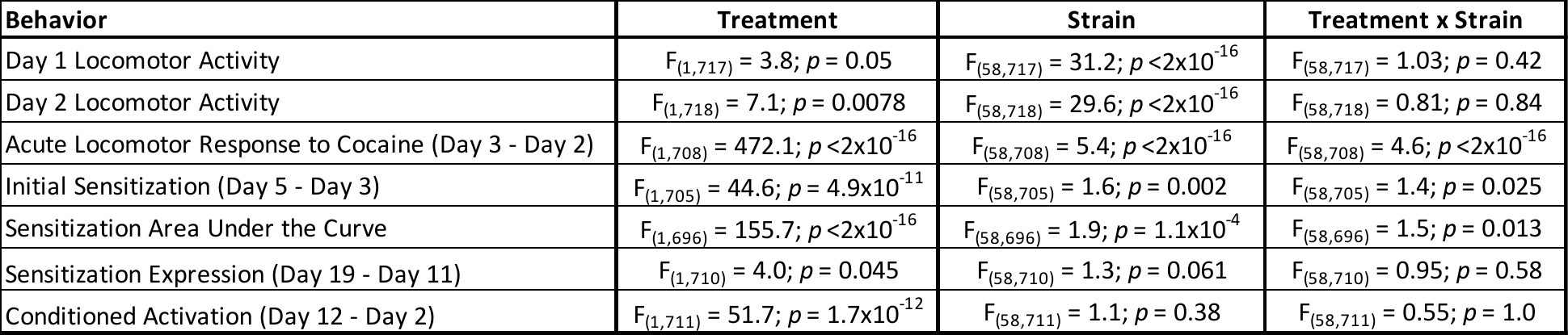

**Supplemental Table 4.** Founder strain sex differences in locomotor response to cocaine

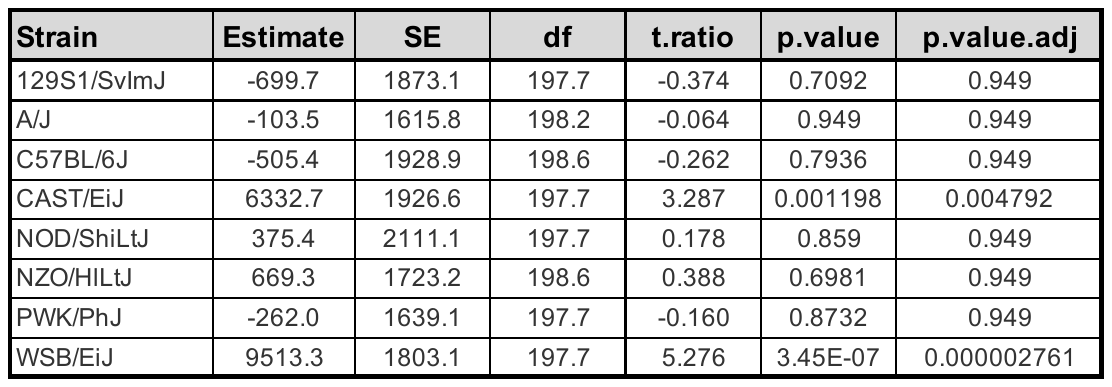

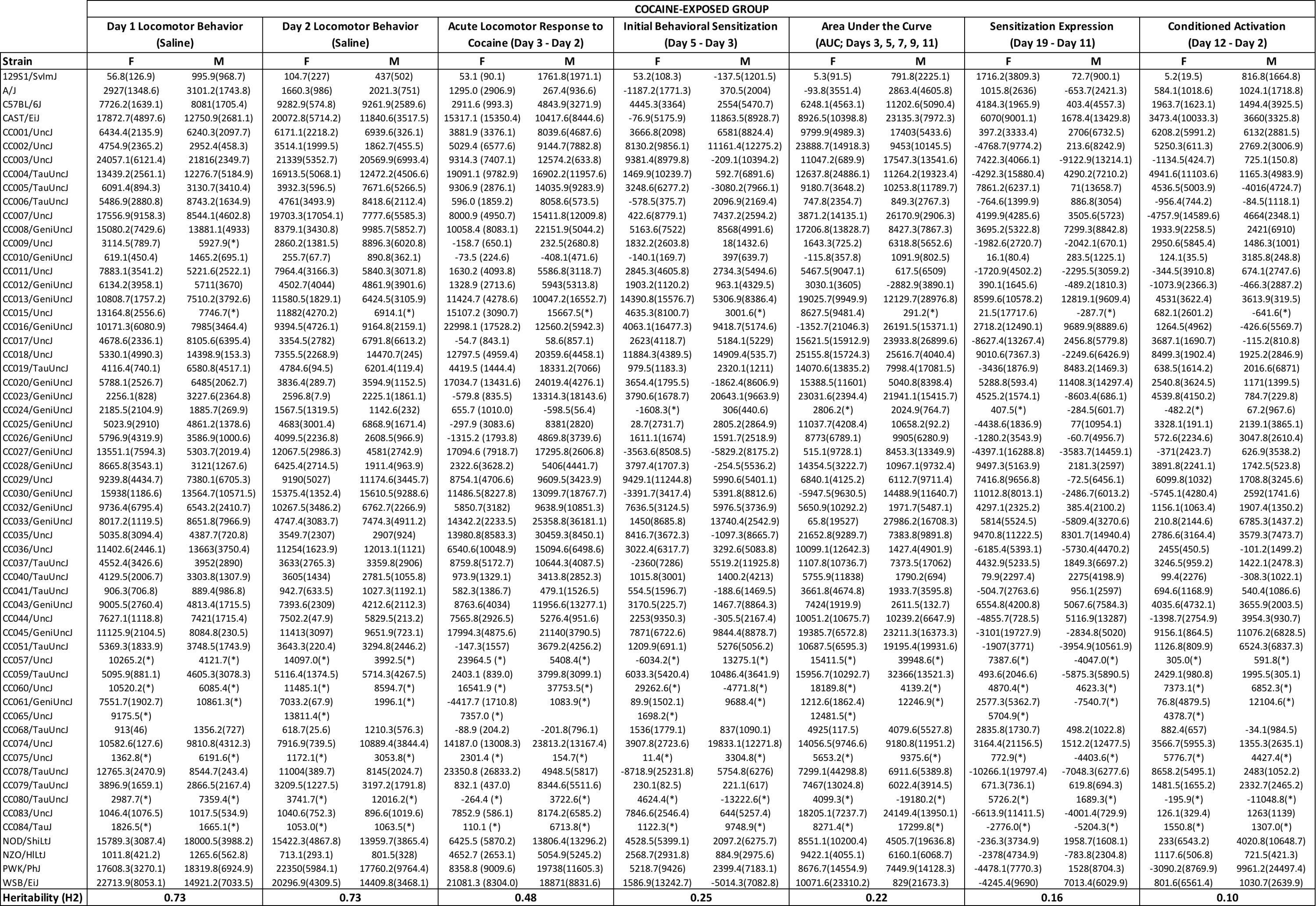

**Supplemental Table 5A.** All strain means (stdev) separated by sex for derived variables in CC and Founder mice exposed to cocaine

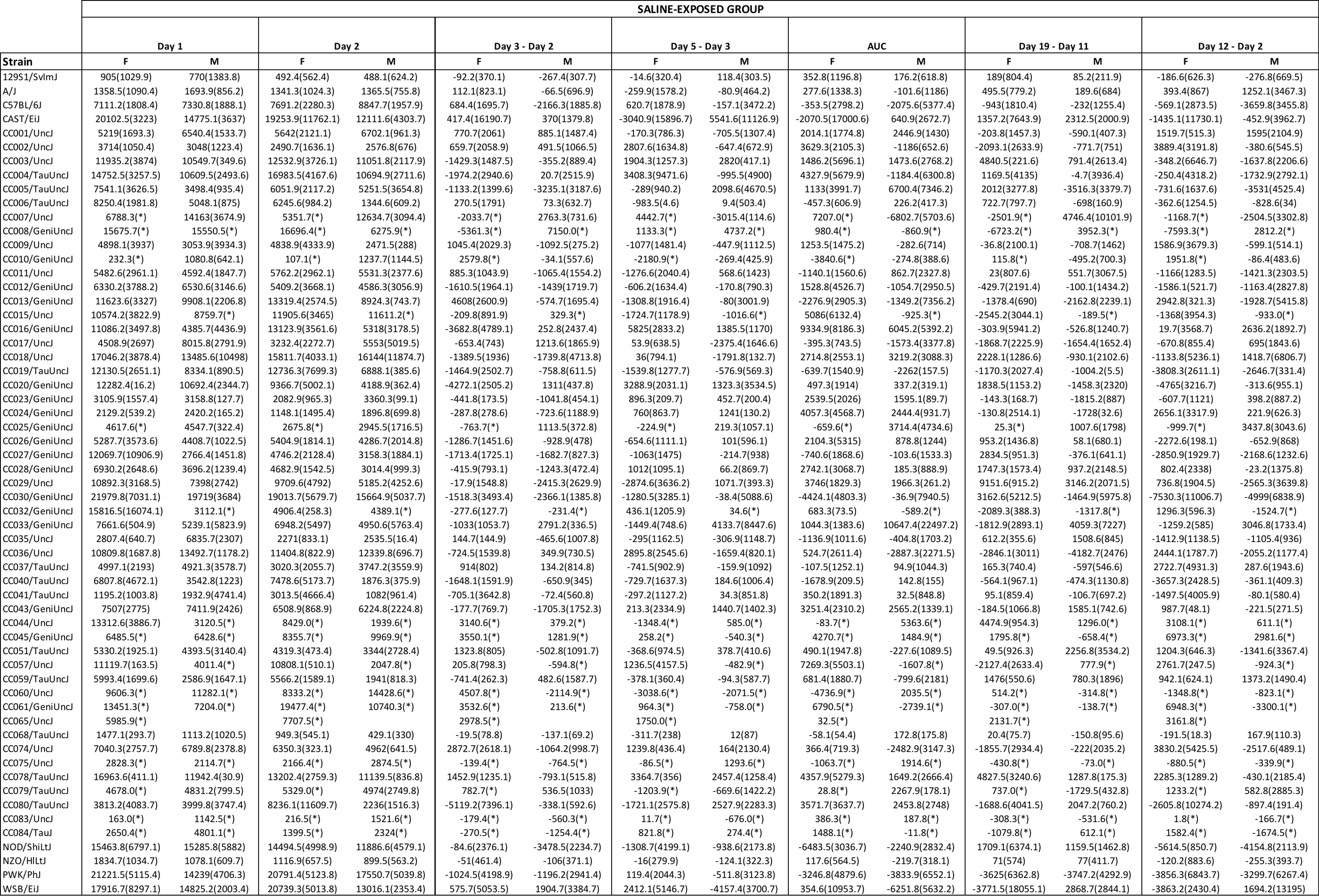

**Supplemental Table 5B.** All strain means (stdev) separated by sex for derived variables in CC and Founder mice exposed to saline

**Supplemental Table 6**. Results of Spearman correlation analysis of strain means for derived variables in Collaborative Cross and Founder strains exposed to cocaine

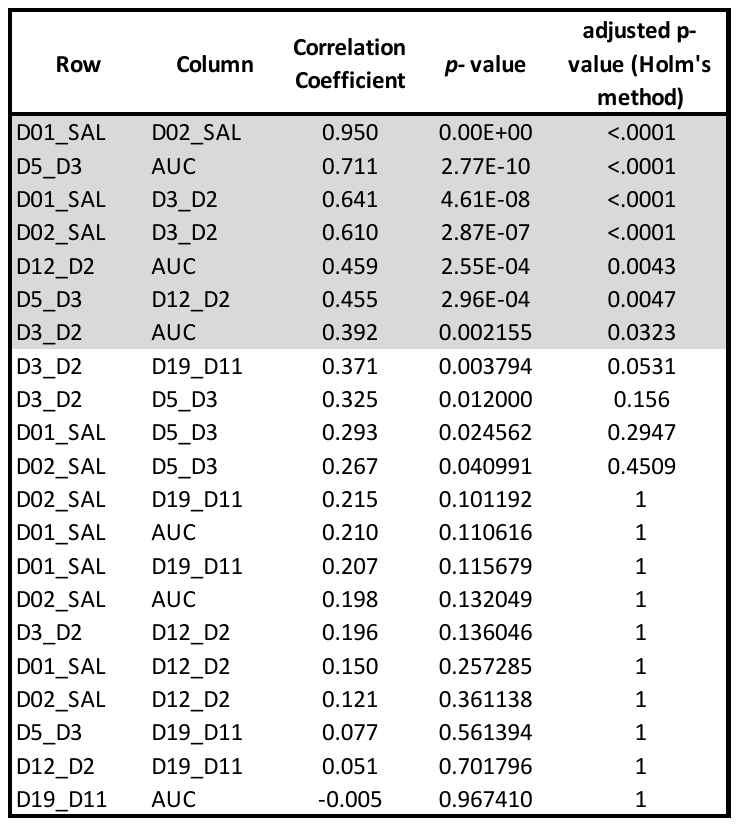

**Supplemental Figure 1.** Collaborative Cross and Founder strain distribution for Day 1 (**A**) and Day 2 (**B**) locomotor activity, acute locomotor response (**C**; Day 3 – Day 2), initial sensitization (**D**; Day 5 – Day 3), sensitization AUC (**E**), sensitization expression (**F**; Day 19 – Day 11) and conditioned activation (**G**; Day 12 – Day 2)

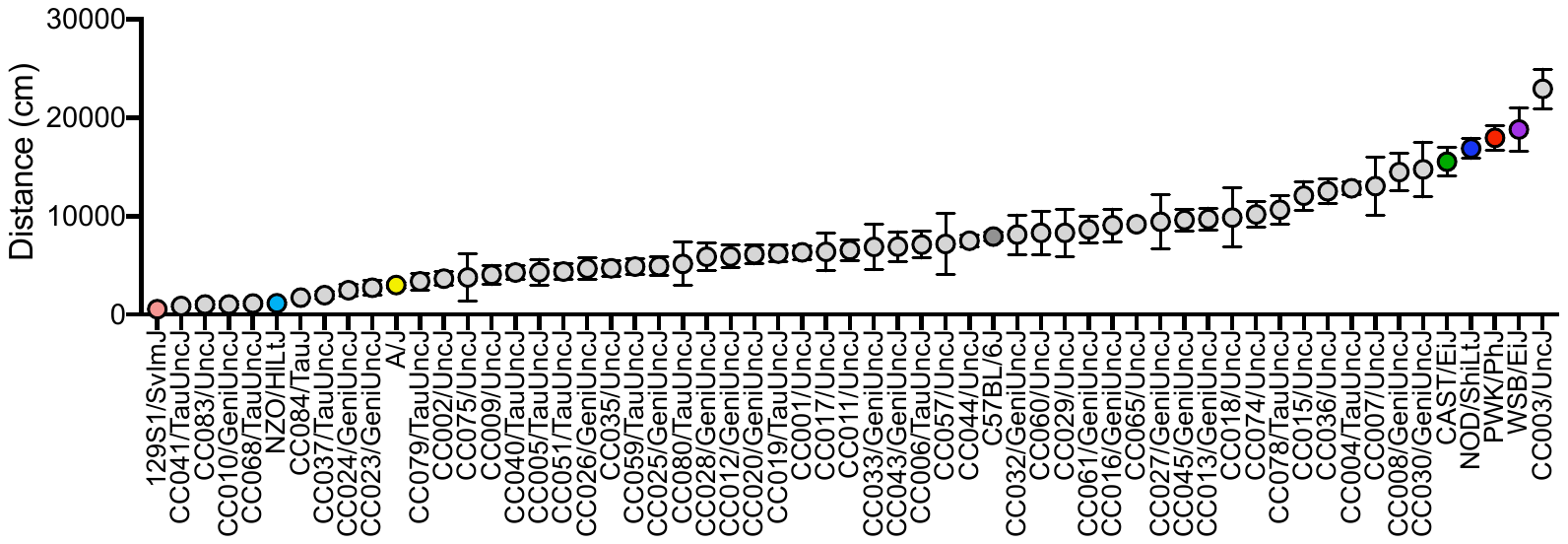

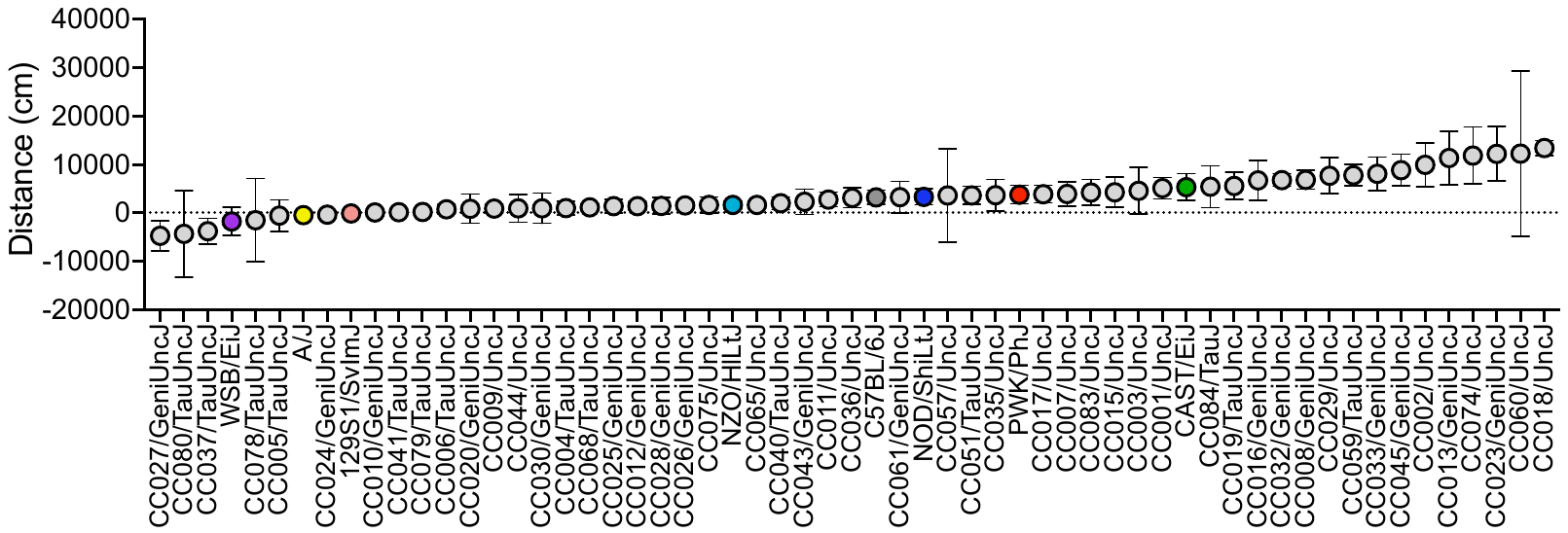

Day 1

Day 5 – Day 3

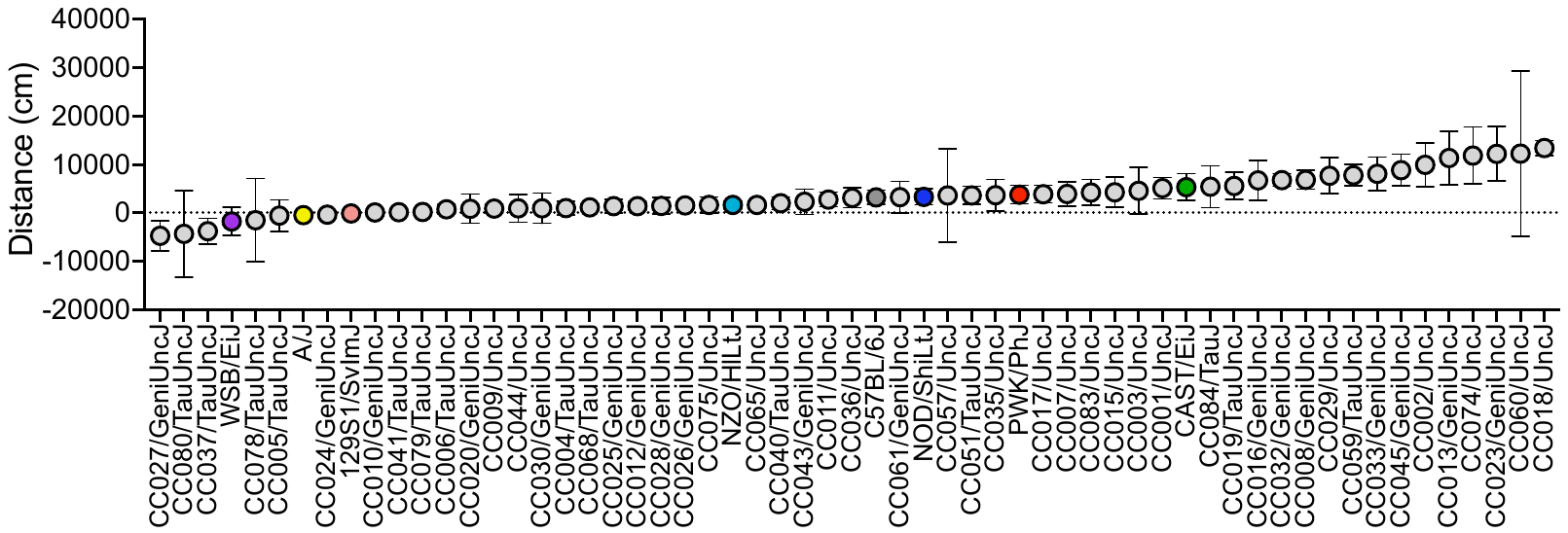

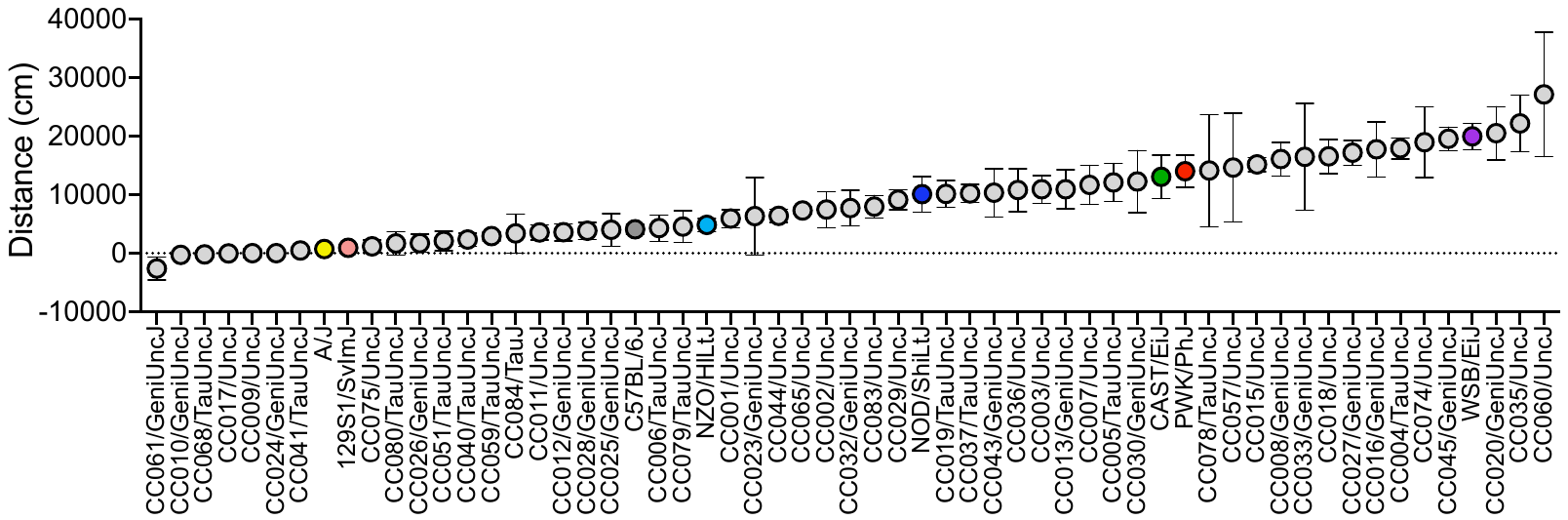

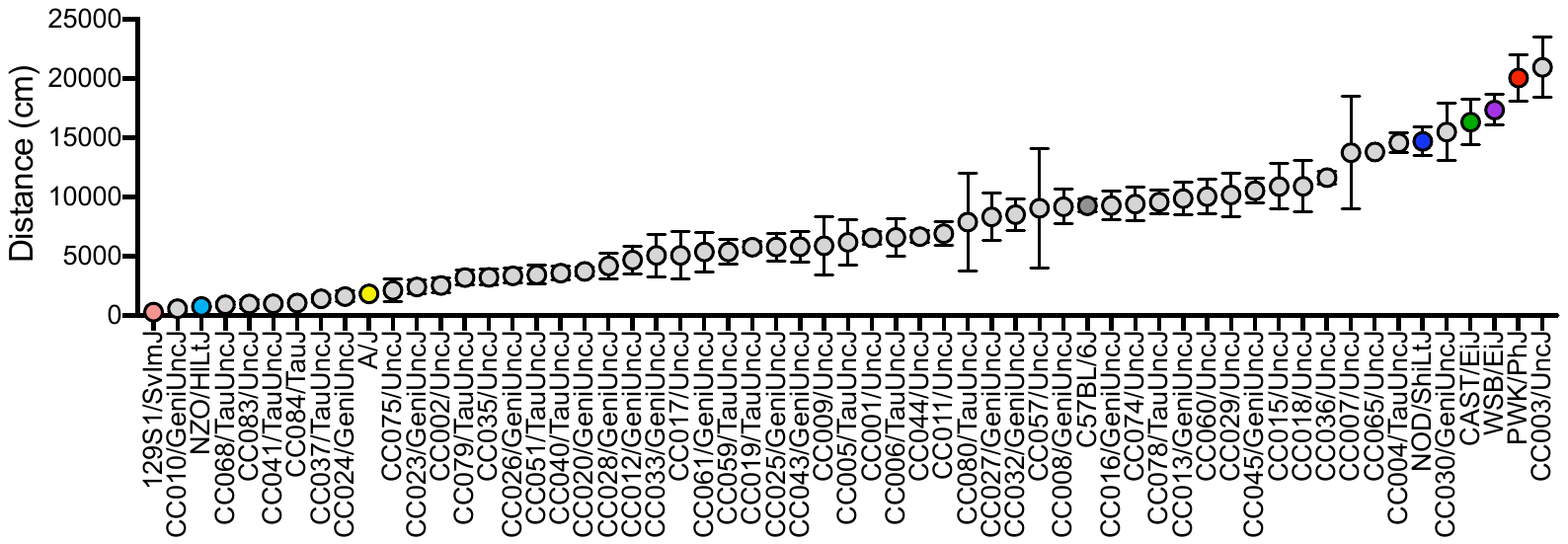

Day 5 – Day 3

D

C

Day 3 – Day 2

Day 2

B

A

**Supplemental Figure 1 cont’d**

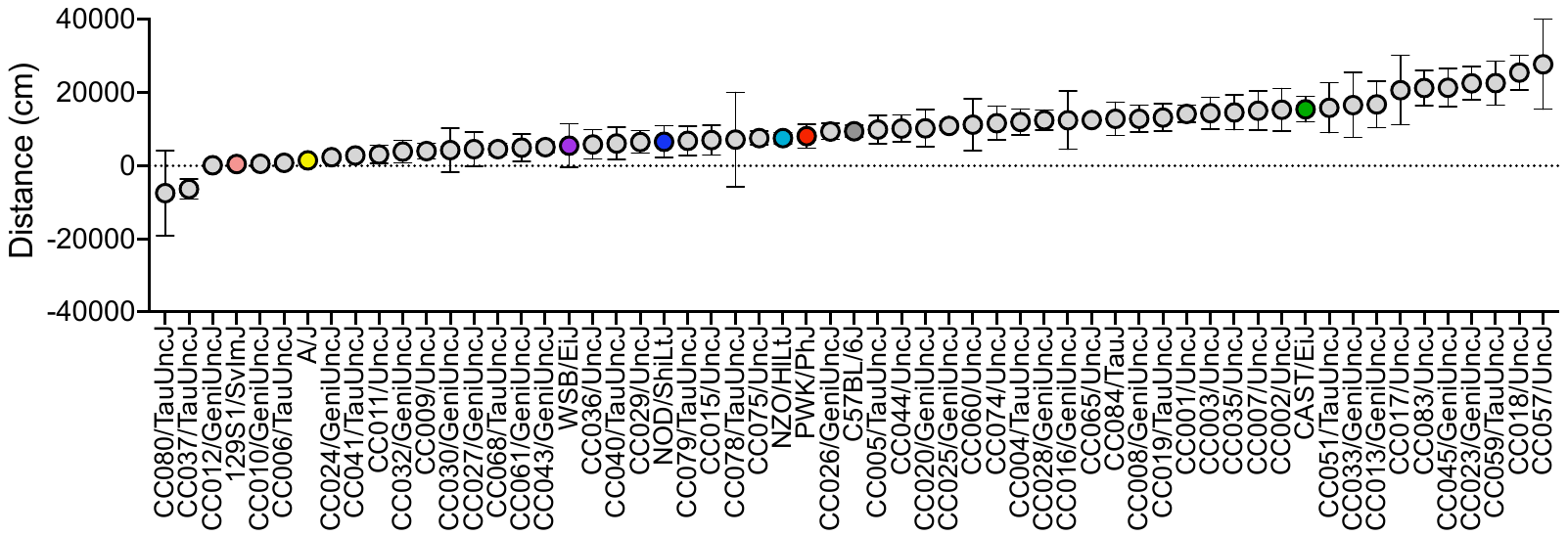

Area Under the Curve

E

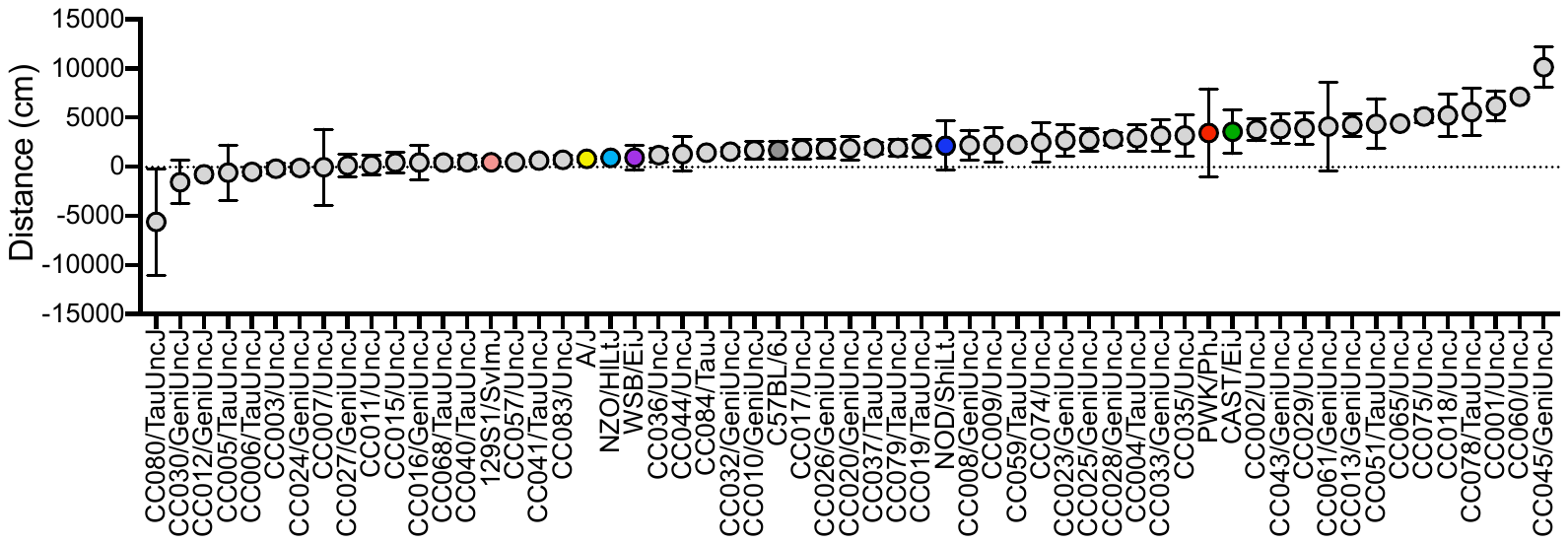

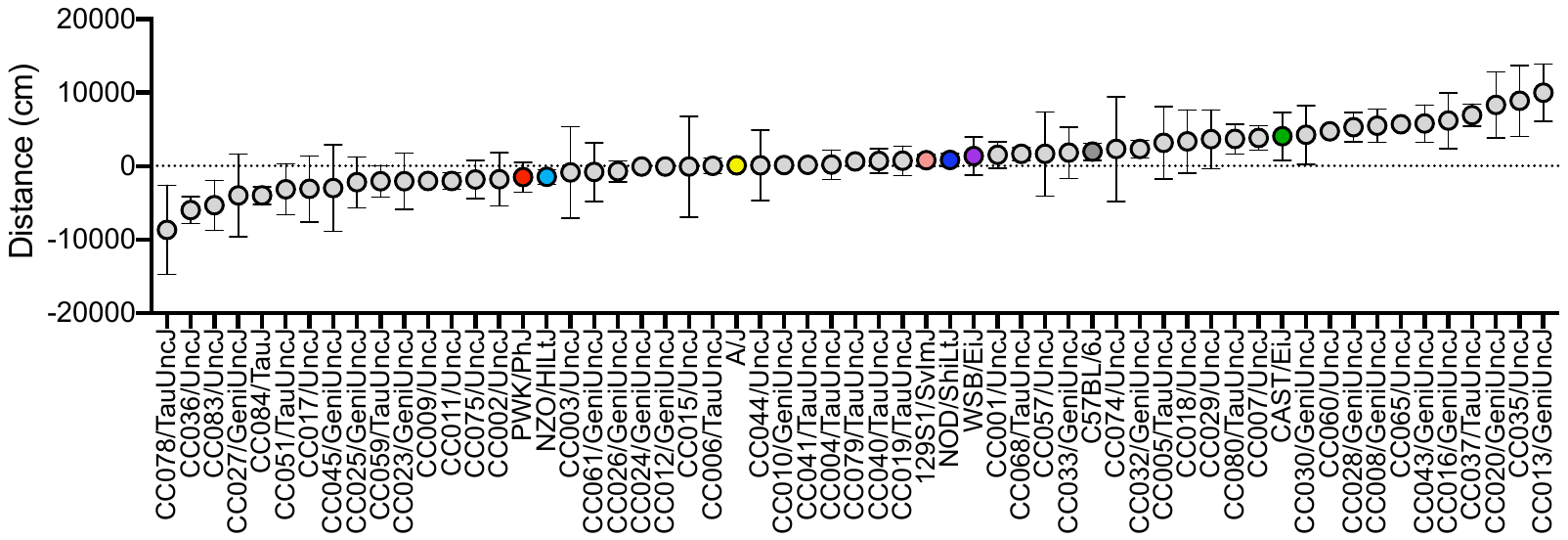

Day 12 – Day 2

G

F

Day 19 – Day 11

**Supplemental Figure 2.** Individual founder strain data separated by sex and treatment

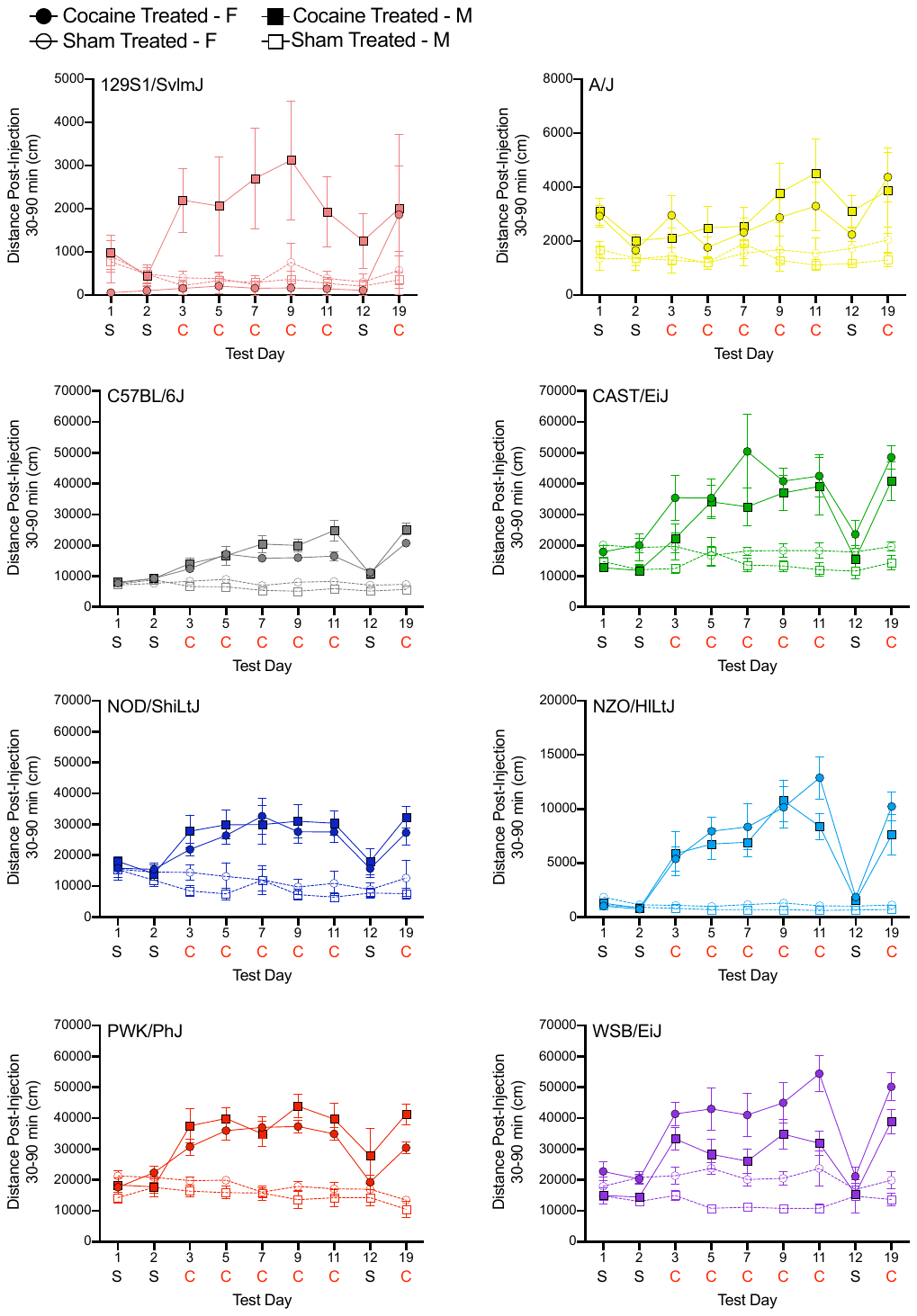

H

G

E

F

D

C

B

A

**Supplemental Figure 3.** Comparison of 19-day behavioral sensitization data for CC016/GeniUncJ (**A**), CC061/GeniUncJ (**B**) and CC074/UncJ (**C**) mice tested at JAX and UNC.

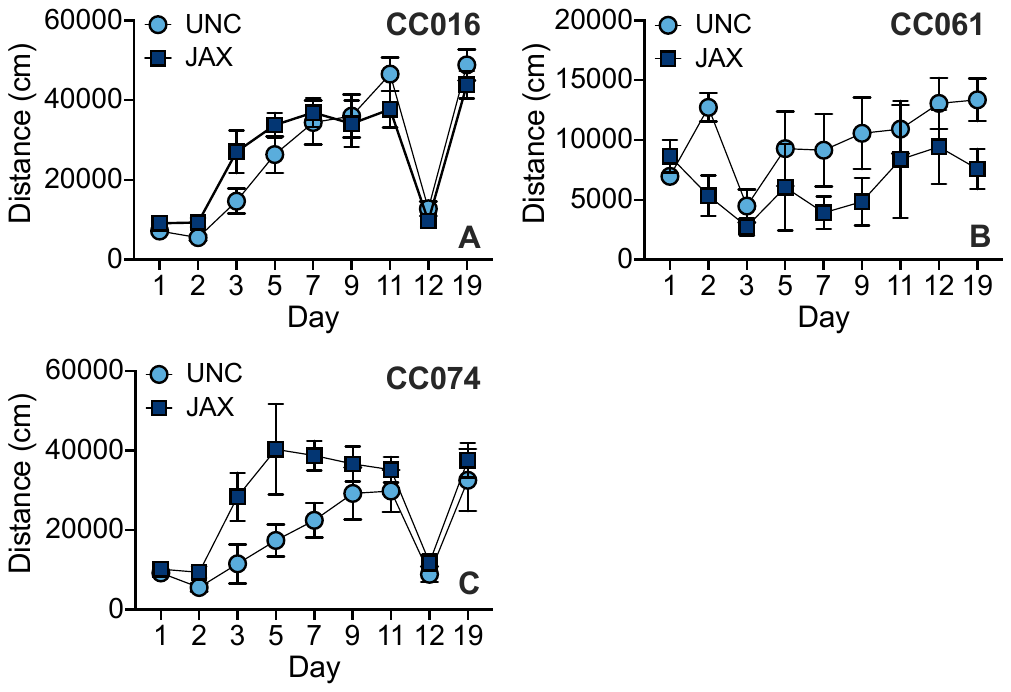
